# Meta-analysis of diets used in *Drosophila* microbiome research and introduction of the *Drosophila* Dietary Composition Calculator (DDCC)

**DOI:** 10.1101/2020.03.17.995936

**Authors:** Danielle N.A. Lesperance, Nichole A. Broderick

## Abstract

While the term standard diet is commonly used in studies using *Drosophila melanogaster*, more often than not these diets are anything but standard, making it difficult to contextualize results in the broader scope of the field. This is especially evident in microbiome studies, despite diet having a pivotal role in microbiome composition and resulting host-microbe interactions. Here, we performed a meta-analysis of diets used in fly microbiome research and provide a web-based tool for researchers to determine the nutritional content of diets of interest. Our goal is for these community resources to aid in contextualizing both past and future microbiome studies (with utility to other fields as well) to better understand how individual lab diets can contribute to observed phenotypes.

## Introduction

In the laboratory, the typical *Drosophila melanogaster* diet is composed of agar, yeast, a sugar source, and cornmeal. However, in reality dietary compositions vary greatly across laboratories, making it difficult to clearly define the composition of a “standard” fly diet. Multiple “branded standard” diets exist such as the Bloomington Standard or CalTech diets that originated at hubs of *D. melanogaster* research, and while many lab groups base their own diets on these recipes, the vast majority of groups maintain flies on diets unique to their laboratory. Differences between these diets, despite their general suitability for fly rearing, can make it challenging to contextualize studies within the scope of *D. melanogaster* research, as nutrition is a critical factor influencing many aspects of physiology including metabolism (Piper *et al.* 2005; Brookheart and Duncan 2016), behavior (Edgecomb *et al.* 1994; Ormerod *et al.* 2017; Davies *et al.* 2018), development (Ormerod *et al.* 2017; Grangeteau *et al.* 2018), longevity (Piper *et al.* 2005; Ormerod *et al.* 2017; Stefana *et al.* 2017), and microbiome composition and function (Wong *et al.* 2014; Obadia *et al.* 2018; Erkosar *et al.* 2018). The relationship between nutrition and the gut microbiome is particularly important, as altering one will likely impact the other with physiologic consequences. Diet plays a pivotal role in shaping microbiome composition and affects interactions between microbiota and host, and the microbiome itself impacts the fly’s nutritional environment, both as a direct source of nourishment and via production and/or utilization of nutrients (Storelli *et al.* 2011; Shin *et al.* 2011; Wong *et al.* 2014; Yamada *et al.* 2015; Huang and Douglas 2015; Broderick 2016; Keebaugh *et al.* 2018; Erkosar *et al.* 2018; Keebaugh *et al.* 2019). Together, dietary nutrition and the microbiome act in concert with one another to dictate nutritional physiology (**Figure 1**).

**Figure 1.**
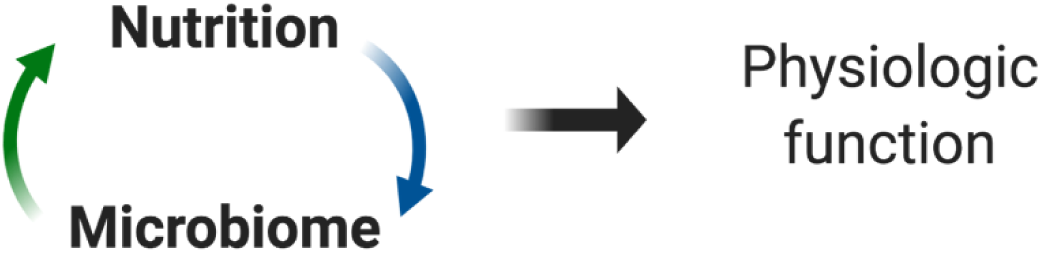
Dietary nutrition and the microbiome are inextricably linked. Dietary nutritional content impacts the diversity and abundance of microbiome members, can influence microbe-microbe interactions, and affects metabolites produced by the microbiome. At the same time, the microbiome itself contributes to overall nutrition via production of metabolites, which are then utilized by the host, catabolism of carbohydrates, and by serving as a direct source of protein to the fly. Together, dietary nutrition and the microbiome interact to play a significant role in host physiology.

In an effort to aid in the contextualization of studies focused on the *D. melanogaster* microbiome, we performed a meta-analysis of diets used across the field. We analyzed the nutrition values of diet recipes, focusing on protein and carbohydrate content of diets to visualize how widely “standard” laboratory diets vary across *D. melanogaster* microbiome studies. Additionally, we have provided a web-based tool for use by the broader community that we’ve named the Drosophila Diet Composition Calculator (DDCC, https://www.brodericklab.com/DDCC.php), which can be used to rapidly determine the macronutrient content of diets of interest simply by inputting amounts of each diet component for a given diet. It is our hope that this meta-analysis and the DDCC can be used to better understand dietary influences on previously observed phenotypes and serve as a resource for experimental design of future studies involving fly nutrition.

## Methods

### Nutritional information for dietary components

Values for calories, fiber, sugars, protein, fat, and carbohydrates were determined for each dietary component using nutritional labels for specific food products, information directly from manufacturers, or from NutritionData.com, a database of food nutritional values obtained from the United States Department of Agriculture’s National Nutrient Database for Standard Reference. The sources for each dietary component are provided in the Supplemental Files. The carbohydrate and protein information for raw fruits was determined using NutritionData.com.

### Analysis of dietary differences across microbiome studies-Fly Microbiome Diet Database

Dietary compositions from over 50 articles (listed in **TableS1**) with a focus on the *D. melanogaster* microbiome were recorded in appropriate columns of the database (Columns A-AF). Calculations for calories per liter, grams of fiber per liter, sugars per liter, protein per liter, fat per liter, carbohydrates per liter, percent fiber, percent sugars, percent protein, percent fat, percent carbohydrates, and the ratio of protein to carbohydrates (P:C) (Columns AH-AT) were performed within the spreadsheet using the previously determined nutritional value for each dietary component. Nutritional information for the holidic fly diet (Piper *et al.* 2014) was determined by inputting the agar and sucrose amounts in the spreadsheet as normal and adding the calculated final mass of amino acids per liter to the formula in Column AL (grams of protein per liter). Similarly, for other diets containing one unique ingredient not otherwise represented in the database, calculations were performed as normal with the nutritional information for the unique ingredient added manually. In these cases, notes are made on the database to indicate special calculations. If it was not possible to calculate the nutritional information for an individual diet, it is noted in Columns AH-AM. Articles that did not readily provide dietary composition were documented for analytical purposes but excluded from the publicly available database. Ultimately, six “branded standard” diets and 71 explicitly reported diets from the literature were included in the database. An additional 14 studies examined did not provide their dietary composition.

### The Drosophila Dietary Composition Calculator (DDCC)

Calculations used to obtain the nutrition facts for the database were used to generate the calculator tool found at https://www.brodericklab.com/DDCC.php. Through this web-tool we also invite researchers to submit published diets using the provided web form to be placed in the publicly available database.

### Data availability

The source files for all nutritional information used to create the Fly Microbiome Diet Database and the DDCC are located at [https://doi.org/10.6084/m9.figshare.11920743.v1]. A downloadable version of the Fly Microbiome Diet Database is located at [https://doi.org/10.6084/m9.figshare.11920788.v2].

## Results and Discussion

### Comparison of diets used across fly microbiome studies

We analyzed the nutritional content of over 70 published diets used for *D. melanogaster* microbiome research based on the dietary components listed in the study methods. Dietary composition varies considerably both in the types of components used and the amounts of components, leading to a wide range of calories, protein, carbohydrate, fat, and fiber levels (**Figure 2**). Moreover, the type/source of a given ingredient can impact these values. For example, for a common ingredient like yeast, several different formulations are used including active, inactive, brewer’s, Lesaffre, and Springaline, all of which have unique nutritional compositions (e.g. protein content ranges from 38% in active dry yeast to 63% in Springaline yeast). Specific ingredients can also add unexpected components to diet. For example, Springaline yeast (BioSpringer), used by a number of European fly immunity/microbiome labs contains 0.03 grams of the antioxidant glutathione per gram of yeast, meaning typical diets can range from 1.5-1.8 grams of added glutathione per liter of diet. This equates to a concentration of around 5 mM, a level used in some studies to block superoxide toxicity (Kim *et al.* 1997; Buchon *et al.* 2009).

**Figure 2.**
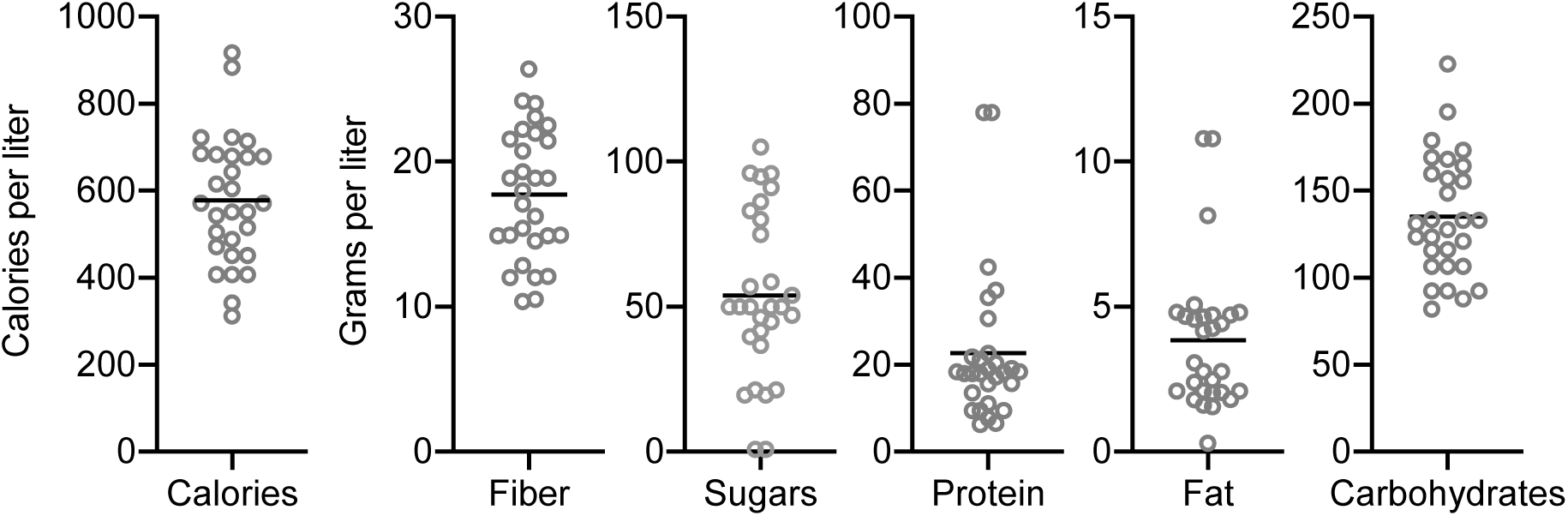
Nutritional content of “standard” *D. melanogaster* diets. Calories, grams of fiber, grams of sugars, grams of protein, grams of fat, and grams of carbohydrates per liter of food of laboratory diets reported as “standard” in the literature. Each point represents a different diet. The minimum and maximum values for each parameter as are follows: Calories-311.97 and 917.13, Fiber-10.36 and 26.38, Sugars-0.80 and 105.00, Protein-6.33 and 77.93, Fat-0.30 and 10.80, Carbohydrates-81.90 and 222.71. Line represents mean. n=29 diets referred to as “standard” out of 71 diets.

To get a better sense for nutritional differences across the diets, we focused on protein and carbohydrate content (**Figure 3A**). While some overlap was seen, particularly for “branded standards” or multiple studies from the same laboratory, the overall spread of protein and carbohydrate content was large. Dietary protein to carbohydrate (P:C) ratio is known to be an important factor influencing life history traits (Lee *et al.* 2008; Jang and Lee 2018), so we next compared P:C of each diet and identified a range of maintenance diets (i.e. not experimental diets with altered diet components) with P:C’s from 0.05 to 0.86 (**Figure 3B**). We additionally noted that a range of P:C’s existed for diets considered “rich” or “poor” with regard to protein content. “Poor” diet P:C’s were between 0.03 and 0.69 with “rich” diets ranging from 0.05 to 0.8 (**Figure 3B**).

**Figure 3.**
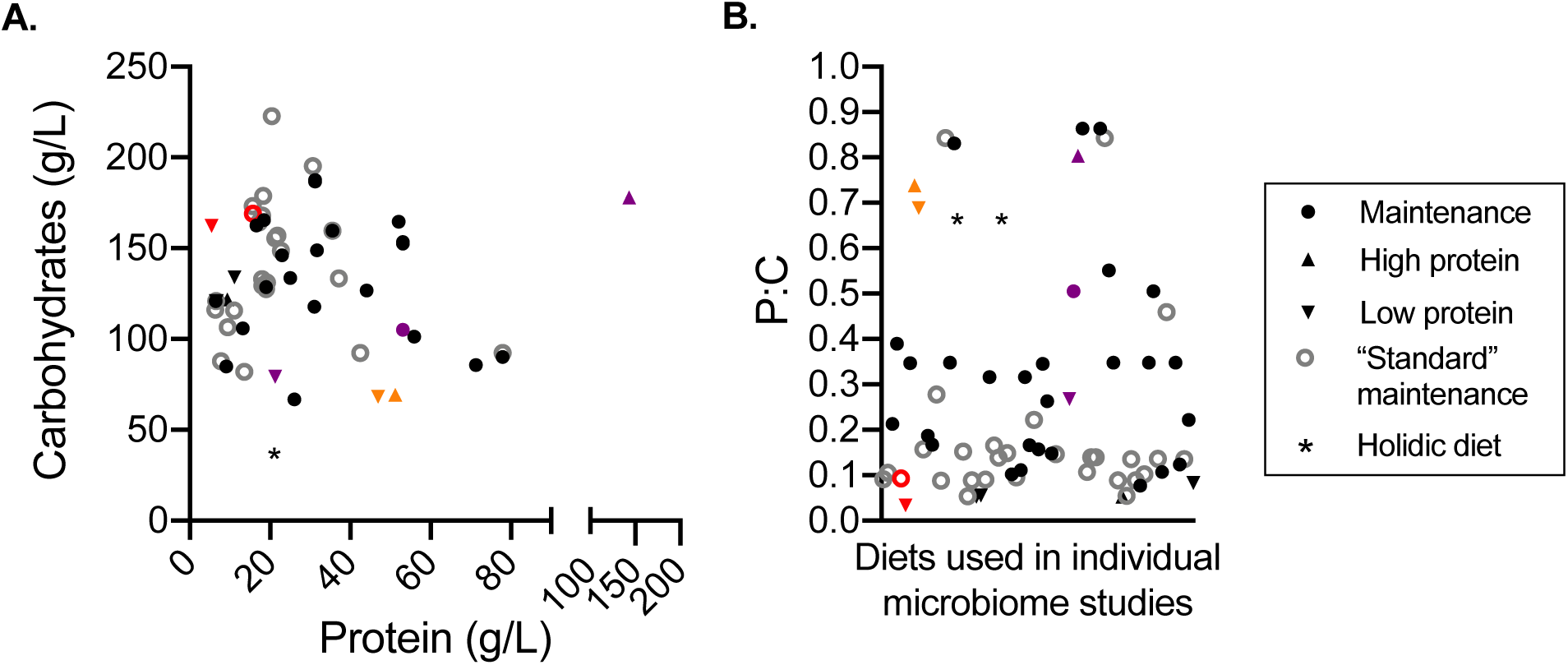
Comparisons of diets used across microbiome research. **A)** Protein and carbohydrate content of diets as determined using the microbiome database. **B)** Protein-to-carbohydrate ratio (protein divided by carbohydrates) of individual diets. Each point represents a different diet reported in fly microbiome literature: closed circles represent diets used for normal maintenance of fly lines; triangles represent diets specifically defined as “rich” or high protein; inverted triangles represent diets designated as “poor” or low protein; open grey circles represent maintenance diets that are described as “standard” in the literature; asterisks represent the holidic fly diet. In **(B)**, red points are examples of two diets used in the same study that represent both a normal and low protein diet (Shin *et al.* 2011); orange points similarly represent another study utilizing a high and low protein (Storelli *et al.* 2011); purple points represent a third study using multiple diets (Erkosar *et al.* 2018). n=71 diets (14 diets were not provided).

Using this visualization of dietary composition, we observed an interesting comparison between two studies that each demonstrated a role for the microbiome in normal larval development in protein poor conditions (achieved through reduced yeast levels; Storelli *et al.* 2011 and Shin *et al.* 2011). Shin *et al.* used two diets that are relatively low in protein (red points) and only differed in P:C by 0.06. Storelli *et al.* also used two diets that differed in P:C by a similar level (0.05), however compared to Shin *et al.* these diets were relatively protein rich (orange points). Both studies show that the microbiome enhanced fly development on their respective low protein diets, but not on the higher protein version. Our comparative analysis indicates that small shifts in protein, even if not evident from P:C values, can be sufficient to reveal biologically important phenotypic effects of diet. However, while the observed phenotypes were similar in these studies, different mechanisms behind the observed developmental effects were reported, including being attributed to different microbiome members-*Acetobacter pomorum* in Shin *et al.* and *Lactobacillus plantarum* in Storelli *et al.* Our analysis shows that the overall diets differ significantly in both protein and carbohydrates levels (**Figure 3**), which could explain the different microbes and mechanisms, as macromolecule concentrations could greatly impact microbiome composition, microbe and/or host physiology, and/or the resulting interaction. This is supported by recent work by Erkosar *et al.* who showed that flies reared on diets containing significantly different concentrations of yeast (**Figure 3**, purple points) had distinct shifts in microbial community composition (Erkosar *et al.* 2018). These examples highlight the importance of contextualizing studies based on dietary composition and how such comparisons can influence interpretation and subsequent studies.

### The “standard diet” fallacy

At the time of writing, 16% of articles examined (14 of 85) gave no clearly defined diet composition and of this group, 71% (10 of 14) described their diet as “standard.” Overall, 46% of diets from all articles (39 of 85) were referred to as “standard,” yet both the range of diet components and total nutritional values of these diets are large (**Figure 2** and shown as open grey in **Figure 3**). It is clear from the ranges we observed that no true “standard” diet exists, highlighting the problematic, but common, phrasing of “standard fly diet” in the literature, which is compounded when the diet recipe is not provided. Our analysis only looked at fly microbiome studies, but we expect this is a wide-spread problem and that other areas of *D. melanogaster* research have a similarly wide range of “standard” diets (whether explicitly reported or not).

### Artificial versus natural diets

To understand how the range of laboratory diets compares to natural fruit diets that *D. melanogaster* encounters in the wild, we obtained protein and carbohydrate information (grams per kilogram) for apples, pears, grapes, bananas, oranges, limes, peaches, and lemons. Carbohydrates spanned from 93 g/kg to 228 g/kg and protein from 3 g/kg to 11 g/kg, resulting in a range of P:C’s from 0.02 to 0.11 (**Figure 4**). While many artificial diets fall within this range, protein content is typically much higher in laboratory conditions compared to natural diets, which may contribute to the lower diversity of microbes found in laboratory reared flies compared to wild-caught (Chandler *et al.* 2011; Erkosar *et al.* 2018). In either natural or artificial diets, however, the nutritional role of microbes must also be considered. In nature, *D. melanogaster* only associates with decomposing (ripe/over-ripe) fruit that support high densities of yeasts and bacteria, which presumably alter macronutrient content of the food while also providing nutrients directly. While artificial diets remove the requirement for microbes to break down complex plant material before consumption by the fly, microbes likely still impact nutrition in artificial diets, but the extent of this and its impacts on the fly in “standard” conditions has not been extensively explored.

**Figure 4.**
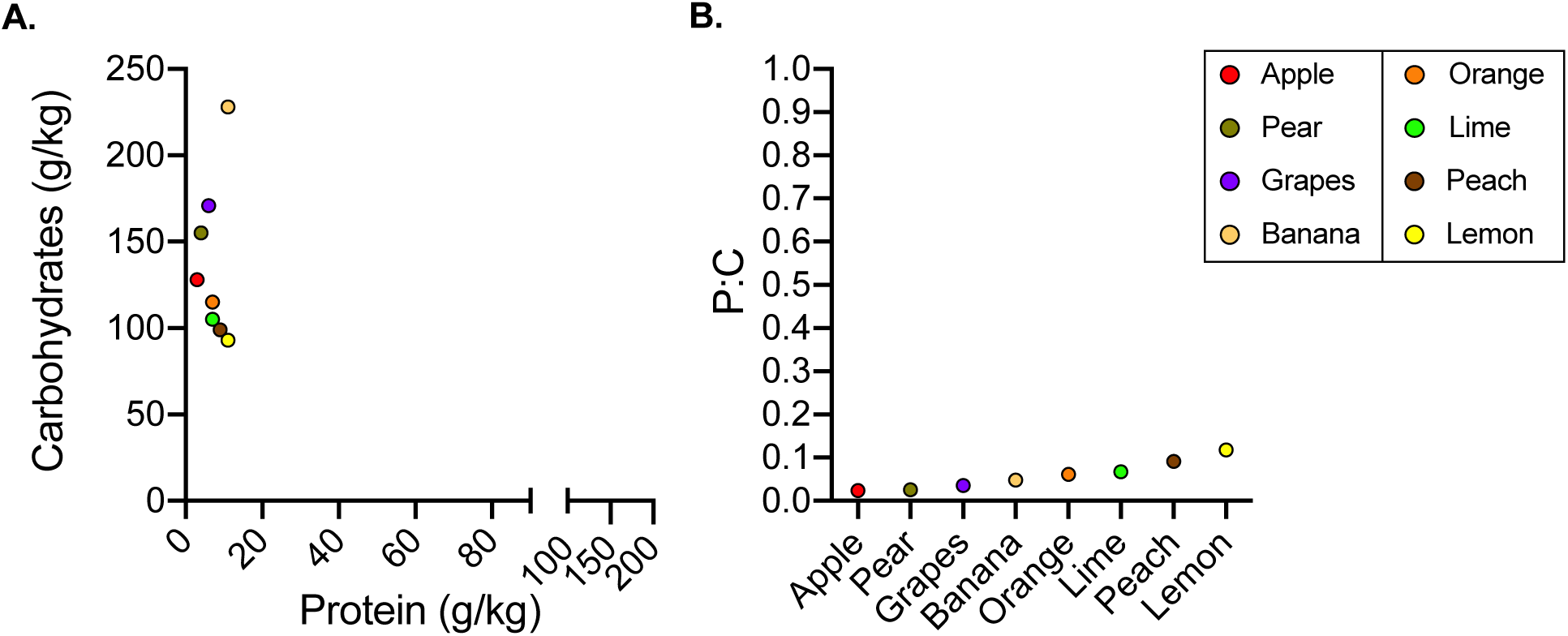
Comparison of protein and carbohydrate content of fruits. **A)** Protein and carbohydrates of raw fruits. **B)** Protein-to-carbohydrate ratio (protein divided by carbohydrates) of raw fruits. Each point represents nutritional information for a different fruit as provided by the United States Department of Agriculture.

### Does D. melanogaster need a standard diet?

It is clear that differences in fly diet have led to issues in reproducibility of results across the field (See Sharon *et al.* 2010, Obadia *et al.* 2018, and Leftwich *et al.* 2018 for one example; Douglas 2018 for commentary on another). One approach to combat such issues is the use of a fully defined diet such as the holidic diet (Piper *et al.* 2014). There are many advantages of using a chemically defined diet, as diet components are more strictly controlled, providing greater power to assess the role of individual diet components on host physiology and microbiome-mediated impacts. However, chemically defined diets are labor-intensive to make and are less representative of natural, complex dietary substrates (which include complex textures, different particle sizes, etc.) making this an unrealistic option for standardization of fly rearing and research across fields. We suggest that a manageable and reasonable approach to address dietary differences across studies is simply to require explicit reporting of diet composition at the time of publication. While having such data does not eliminate variability, it is invaluable for contextualizing results and phenotypes, provides potential explanations for observed differences, and testable hypotheses for follow-up in subsequent studies. We also expect that use of complex diet components is beneficial for discovery of physiologically relevant phenotypes that may otherwise be lost or artificially altered on more defined diets. For example, food particle size in animal gut ecosystems is known to impact digestion and bulk passage rate as well as microbiome composition through attachment and microcolony support (Cheng *et al.* 1981; Martz and Belyea 1986; Bjorndal *et al.* 1990; McAllister *et al.* 1994; Vermeulen *et al.* 2018; Kiarie and Mills 2019). Ultimately, what is important is that researchers understand the nutritional implications of the diets they use and look to nutritional information as a resource to aid in analysis of results and comparison across laboratories. It is our hope that the examples highlighted in this meta-analysis and the data provided by the DDCC will aid in a broader appreciation for the importance of dietary reporting, and help to contextualize observations across research studies using *D. melanogaster*.

## Web resources

### Fly Microbiome Diet Database

https://doi.org/10.6084/m9.figshare.11920788.v2

### Drosophila Dietary Composition Calculator

https://www.brodericklab.com/DDCC.php

## Acknowledgements

The *Drosophila* Dietary Composition Calculator was created by Big Rose Web Design, LLC. This work was supported by the National Institutes of Health [R35GM128871] and the University of Connecticut.

## Footnotes

None.

## Supplemental data

*Diet component nutritional content reference files FileS1-S21:*

https://doi.org/10.6084/m9.figshare.11920743.v1

---

**Table S1.** Studies referenced in the Fly Microbiome Diet Database.

| DATABASE ROW | CITATION | DOI |
| --- | --- | --- |
| 8 | Brummel *et al.* 2004 | 10.1073/pnas.0405207101 |
| 9 | Ren *et al.* 2007 | 10.1016/j.cmet.2007.06.006 |
| 10, 11 | Sharon *et al.* 2010 | 10.1073/pnas.1009906107 |
| 12, 13 | Shin *et al.* 2011 | 10.1126/science.1212782 |
| 14 | Wong *et al.* 2011 | 10.1111/j.1462-2920.2011.02511.x |
| 15, 16 | Storelli *et al.* 2011 | 10.1016/j.cmet.2011.07.012 |
| 17 | Chandler *et al.* 2011 | 10.1371/journal.pgen.1002272 |
| 18 | Ridley *et al.* 2012 | 10.1371/journal.pone.0036765 |
| 19 | Blum *et al.* 2013 | 10.1128/mBio.00860-13 |
| 20 | Fink *et al.* 2013 | 10.1128/AEM.01903-13 |
| 21 | Staubach *et al.* 2013 | 10.1371/journal.pone.0070749 |
| 22 | Erkosar *et al.* 2014 | 10.1371/journal.pone.0094729 |
| 23 | Newell and Douglas 2014 | 10.1128/AEM.02742-13 |
| 24 | Broderick *et al.* 2014 | 10.1128/mBio.01117-14 |
| 25 | Piper *et al.* 2014 | 10.1038/nmeth.2731 |
| 26 | Clark *et al.* 2015 | 10.1016/j.celrep.2015.08.004 |
| 27 | Deshpande *et al.* 2015 | 10.3945/jn.115.222380 |
| 28, 29, 30 | Yamada *et al.* 2015 | 10.1016/j.celrep.2015.01.018 |
| 31 | Sansone *et al.* 2015 | 10.1016/j.chom.2015.10.010 |
| 32 | Huang and Douglas 2015 | 10.1098/rsbl.2015.0469 |
| 33 | Chaplinska *et al.* 2016 | 10.1371/journal.pone.0167726 |
| 34 | Li *et al.* 2016 | 10.1016/j.chom.2016.01.008 |
| 35 | Galenza *et al.* 2016 | 10.1242/bio.015016 |
| 36 | Sekihara *et al.* 2016 | 10.1074/jbc.M116.761791 |
| 37 | Shanbhag *et al.* 2016 | 10.1113/JP272617 |
| **38** | Elgart *et al.* 2016 | 10.1038/ncomms11280 |
| **39** | Sebald *et al.* 2016 | 10.1371/journal.pone.0153476 |
| **40** | Dobson *et al.* 2016 | 10.1186/s12864-016-3307-9 |
| **41** | Fischer *et al.* 2017 | 10.7554/eLife.18855 |
| **42** | Han *et al.* 2017 | 10.1007/s00248-016-0925-3 |
| **43** | Leitão-Gonçalves *et al.* 2017 | 10.1371/journal.pbio.2000862 |
| **44** | Wong *et al.* 2017 | 10.1016/j.cub.2017.07.022 |
| **45** | Winans *et al.* 2017 | 10.1111/mec.14232 |
| **46** | Obadia *et al.* 2017 | 10.1016/j.cub.2017.05.034 |
| **47, 48, 49** | Jehrke *et al.* 2018 | 10.1038/s41598-018-24542-5 |
| **50, 51, 52** | Erkosar *et al.* 2018 | 10.1002/ece3.4444 |
| **53** | Martino *et al.* 2018 | 10.1016/j.chom.2018.06.001 |
| **54** | Pais *et al.* 2018 | 10.1371/journal.pbio.2005710 |
| **55** | Fast, Duggal, *et al.* 2018 | 10.1128/mBio.01114-18 |
| **56** | Fast, Kostiuk, *et al.* 2018 | 10.1073/pnas.1802165115 |
| **57** | Storelli *et al.* 2018 | 10.1016/j.cmet.2017.11.011 |
| **58, 59** | Téfit *et al.* 2018 | 10.1016/j.jinsphys.2017.09.003 |
| **60** | Inamine *et al.* 2018 | 10.1128/mBio.01453-17 |
| **61, 62** | Keebaugh *et al.* 2018 | 10.1016/j.isci.2018.06.004 |
| **63** | Obata *et al.* 2018 | 10.1038/s41467-018-03070-w |
| **64** | Gould *et al.* 2018 | 10.1073/pnas.1809349115 |
| **65, 66** | Keebaugh *et al.* 2019 | 10.1128/mBio.00885-19 |
| **67** | Su *et al.* 2019 | 10.1186/s12866-019-1686-1 |
| **68** | Solomon *et al.* 2019 | 10.7717/peerj.8097 |
| **69** | Si *et al.* 2019 | 10.1093/gerona/gly189 |
| **70** | Dong *et al.* 2019 | 10.3390/nu11030636 |
| **71** | Tang *et al.* 2019 | 10.1016/j.jinsphys.2019.103893 |
| **72** | Henry *et al.* 2020 | 10.1016/j.cbpa.2019.110626 |
| **73** | Slankster *et al.* 2019 | 10.1893/0005-3155-90.4.227 |
| **74, 75** | Adair *et al.* 2020 | 10.1038/s41396-019-0532-7 |
| **76** | Fast *et al.* 2020 | 10.1016/j.celrep.2019.12.094 |
| **77, 78** | Lee *et al.* 2020 | 10.1007/s00248-019-01404-9 |

